# Towards a global investigation of transcriptomic signatures through co-expression networks and pathway knowledge for the identification of disease mechanisms

**DOI:** 10.1101/2021.03.02.433520

**Authors:** Rebeca Queiroz Figueiredo, Tamara Raschka, Alpha Tom Kodamullil, Martin Hofmann-Apitius, Sarah Mubeen, Daniel Domingo-Fernández

## Abstract

In this work, we attempt to address a key question in the joint analysis of transcriptomic data: can we correlate the patterns we observe in transcriptomic datasets to known molecular interactions and pathway knowledge to broaden our understanding of disease pathophysiology? We present a systematic approach that sheds light on the patterns observed in hundreds of transcriptomic datasets from over sixty indications by using pathways and molecular interactions as a template. Our analysis employs transcriptomic datasets to construct dozens of disease specific co-expression networks, alongside a human interactome network of protein-protein interactions described in the literature. Leveraging the interoperability between these two network templates, we explore patterns both common and particular to these diseases on three different levels. Firstly, at the node-level, we identify the most and least common proteins in these diseases and evaluate their consistency against the interactome as a proxy for their prevalence in the scientific literature. Secondly, we overlay both network templates to analyze common correlations and interactions across diseases at the edge-level. Thirdly, we explore the similarity between patterns observed at the disease level and pathway knowledge to identify pathway signatures associated with specific diseases and indication areas. Finally, we present a case scenario in the context of schizophrenia, where we show how our approach can be used to investigate disease pathophysiology.

## 1. Introduction

Despite the exponential growth of biomedical data in the last decades, we are still far from understanding the function of every gene in a living organism. Nevertheless, major technological advancements now enable us to assign specific biological functions to thousands of protein-coding genes in the human genome (UniProt Consortium, 2019). In turn, complex interactions between groups of genes, proteins, and other biomolecules give rise to the normal functioning of the cell. By acquiring knowledge of these interactions, we can decipher the molecular mechanisms which cause system-wide failures that can lead to disease (Caldera *et al*., 2017). A common modeling approach to represent these vast sets of interactions is in reconstructing mechanisms in the form of networks as intuitive representations of biology, where nodes denote biological entities and edges their interactions (Franzese *et al*., 2019; Winterbach *et al*., 2013).

Numerous standardized formats have been widely adopted to model biological networks that represent pathway knowledge dispersed throughout the scientific literature (Hanspers *et al*., 2020). Pathway models in a variety of formats can be found housed in databases such as KEGG (Kanehisa *et al*., 2021) and Reactome (Jassal *et al*., 2020), each with a varied focus and scope. These databases can be specifically leveraged for hypothesis generation, the analysis of biomedical data such as with pathway enrichment (Reimand *et al*., 2019), or predictive modeling (Segura-Lepe *et al*., 2019). Using the networks of known molecular interactions, one can also discern novel genes involved in particular disease states as functions of network proximity (Huang *et al*., 2018). A general trend noted by Huang and colleagues was the observation that larger networks tended to outperform smaller ones, an effect also observed when comparing the performance of integrated pathway databases to individual ones in enrichment and predictive modeling tasks (Mubeen *et al*., 2019).

Although knowledge-driven approaches that leverage literature-based evidence can be used to gain a mechanistic understanding of disease pathophysiology, these approaches tend to be augmented when applied in combination with data-driven ones. In the latter case, transcriptomic profiling offers researchers a systematic and affordable method to analyze the expression and activity of genes and proteins on a large-scale under distinct physiological conditions. Through gene expression profiling, patterns of genes expressed at the transcript level that are relevant to a particular condition can be determined, whilst considering sets of genes involved in a specific biological process tend to exhibit similar patterns of expression or activity (van Dam *et al*., 2018). To model these patterns, techniques such as gene co-expression networks have been developed in which genes with correlated expression activity are connected. Although several methodologies exist to generate co-expression networks, such as WGCNA (Langfelder *et al*., 2008), they tend to be represented as undirected weighted graphs, where graph nodes correspond to genes, and edges between nodes correspond to co-expression relationships (Stuart *et al*., 2003). The applications of these networks are diverse, ranging from identifying functional and disease-specific modules to hub genes (van Dam *et al*., 2018). For instance, Chou *et al*. (2014) and Xiang *et al*. (2018) combined independent datasets related to endometrial cancer and Alzheimer’s disease, respectively, in order to generate co-expression networks that captured gene expression patterns across multiple disease-specific datasets. Using these co-expression networks, they were able to identify relevant genes in the context of these two indications.

Though it is standard practice to perform enrichment analysis using pathway and gene set databases (e.g., KEGG and Gene Ontology; Gene Ontology Consortium *et al*., 2019) on gene lists from co-expression networks such as those from a particular disease module (Mao *et al*., 2020; Yao *et al*., 2019) for mechanistic insights, this approach ignores the topology of the network as it exclusively relies upon sets of genes rather than the network structure. In a recent study, Paci *et al*. (2021) overcame this challenge by showing how distinct, topological properties of disease networks can emerge through the identification and mapping of disease-specific genes of several disease co-expression networks to a human interactome network of protein-protein interactions.

The potential insights that can be gained from the previously mentioned analyses together with the abundance of publically available transcriptomic datasets (Athar *et al*., 2018; Edgar *et al*., 2002) have prompted the creation of databases that store collections of co-expression networks, such as COXPRESdb for numerous species (Obayashi *et al*., 2019). By harmonizing and storing thousands of transcriptomic datasets in the form of co-expression networks, these resources capture a variety of “snapshots’’ representing gene expression patterns in a diverse set of contexts, such as disease states. With these transcriptomic data and pathway resources in hand, we can connect the transcriptome with the proteome by overlaying the patterns in co-expression networks with the scaffold of biological knowledge embedded in pathway networks. Although such integrative approaches can have multiple applications, they are likely of most relevance for the investigation of disease pathophysiology as systematically combining transcriptomic data with pathway knowledge could reveal insights on specific or shared molecular mechanisms across multiple indications.

In this work, we jointly leverage the patterns of disease-specific datasets reflected in co-expression networks and pathway and interaction networks to uncover the mechanisms underlying disease pathophysiology. To do so, we systematically compared hundreds of transcriptomic datasets from over 60 diseases with a human protein-protein interactome network to unravel the proteins, subgraphs, and pathways that are specific to certain diseases or shared across multiple. Finally, in a case scenario, we demonstrate how bringing together a disease-specific co-expression network with pathway knowledge allows us to better understand the role of a specific pathway within a disease context.

## 2. Methodology

In subsection 2.1, we outline the process of generating disease-specific co-expression networks from transcriptomic data **(Figure 1; left)**. Then, in subsection 2.2, we describe the construction of a human protein-protein interactome network **(Figure 1; right)**. Finally, in 2.3, we outline the various analyses conducted **(Figure 1; center)**.

**Figure 1.**
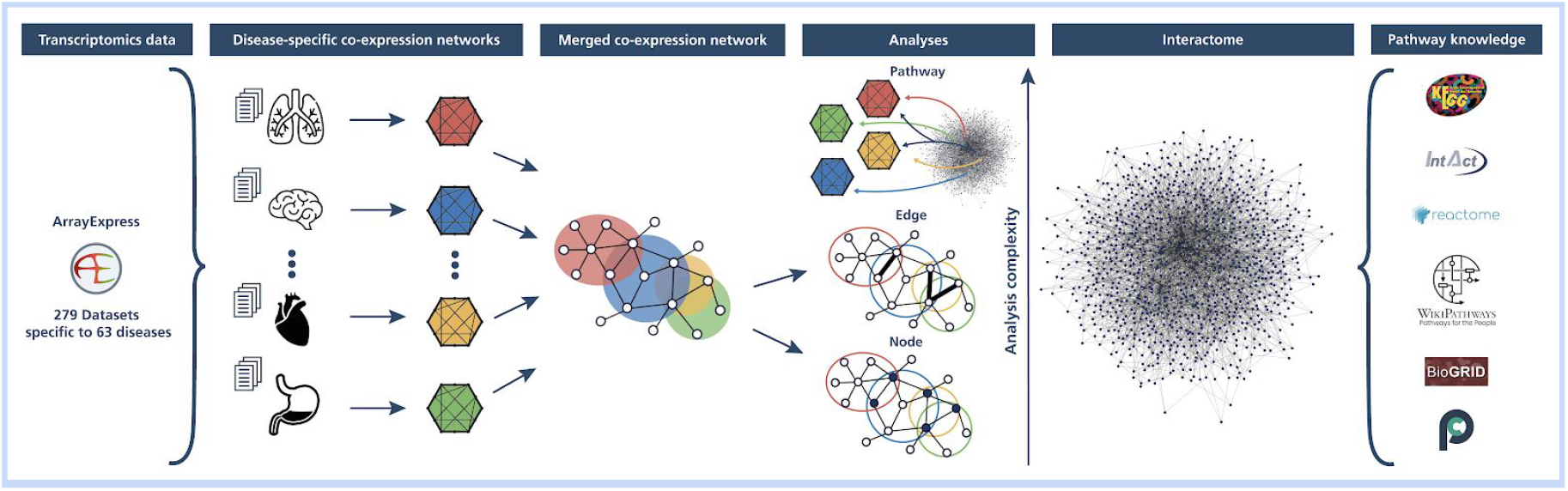
Schematic illustration of the methodology. 279 transcriptomic datasets were acquired from ArrayExpress and grouped into 63 distinct diseases to generate disease-specific co-expression networks **(left)**. A comprehensive protein-protein interactome network was built from an ensemble of six pathway and interaction databases **(right)**. A series of analyses were then conducted on the disease-specific co-expression networks (**center)**, specifically: a node-level analysis **(lower)**, an edge-level analysis **(middle)**, and a pathway-based analysis **(upper)** leveraging pathway knowledge and the interactome network.

### 2.1. Generating co-expression networks from transcriptomic data

#### 2.1.1. Identifying disease-specific datasets in ArrayExpress

We queried datasets from ArrayExpress (AE) (Athar *et al*., 2018) belonging to the most widely used platform: the Affymetrix Human Genome U133 Plus 2.0 Array (accession on AE: A-AFFY-44). By using the same platform for each of the datasets, we ensured that the datasets were relatively comparable. ArrayExpress was preferred over other databases such as Gene Expression Omnibus (GEO) (Edgar *et al*., 2002) as datasets often comprise of normalized and mapped terms in their metadata that describe their characteristics (e.g., experimental details, organism information, etc.). Furthermore, it provides a user-friendly API through which all necessary information was queried. As of 20/07/2020, 4,485 datasets generated from platform A-AFFY-44 have been stored in ArrayExpress, resulting in roughly below 200,000 samples. **Figure 2** summarizes the filtering steps that we conducted to identify disease-specific datasets which are also described below.

**Figure 2.**
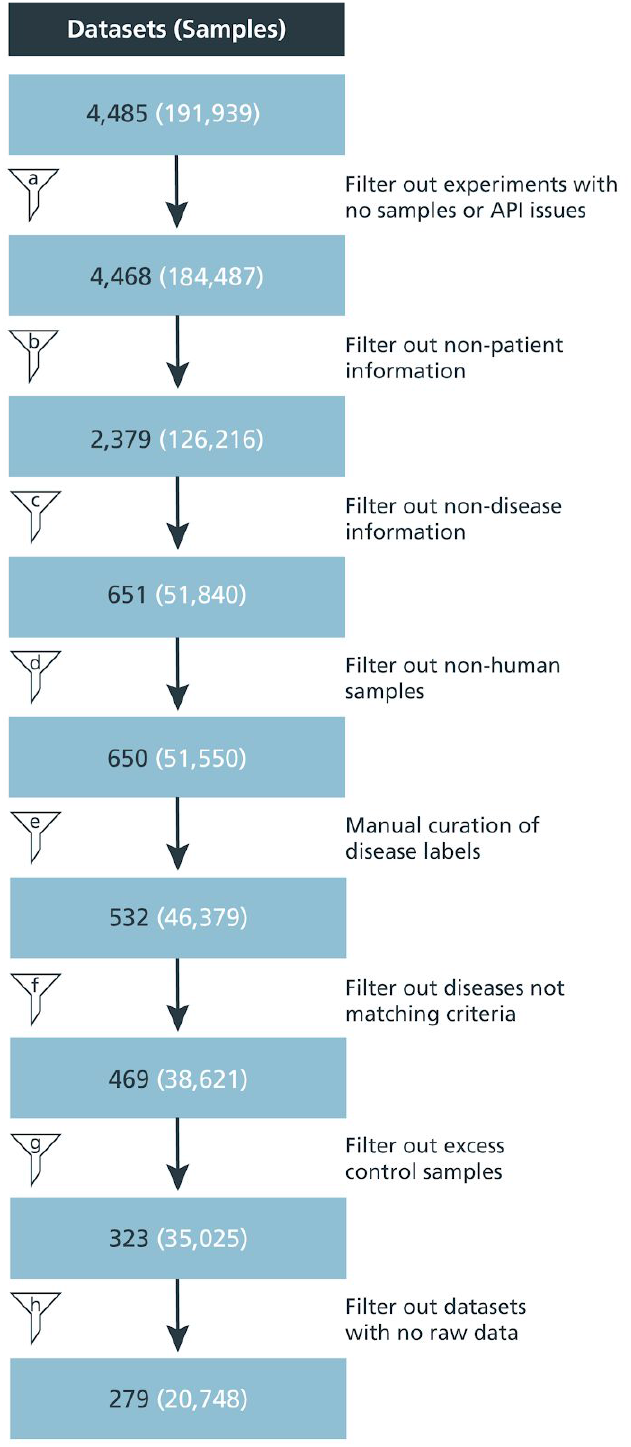
Extracting disease-specific datasets from ArrayExpress. transcriptomic data from nearly 4,500 datasets was derived from ArrayExpress. Several filtering steps **(a-h)** were applied to only retain disease-specific datasets for patient samples and controls that fulfilled the criteria outlined in subsection 2.1.1.

As the purpose of this work was to analyze disease-specific datasets, only patient samples and their controls were eligible for the analysis. Thus, a filtering step was introduced to focus exclusively on patient-level data **(Figure 2, filter A)**. To filter out irrelevant datasets, we leveraged keywords present in the metadata such as ‘dose’, ‘compound’ or ‘strain’ **(Figure 2, filter B)**. Furthermore, information about the disease state of each sample is needed for building disease-specific networks. Therefore, the metadata columns were searched for disease keywords like ‘disease’, ‘histology’, or ‘status’ **(Figure 2, filter C)**. This resulted in 651 datasets, of which one non-human dataset was removed, totaling in 650 datasets with 51,550 samples **(Figure 2, filter D)**.

Once datasets were filtered to identify those that contained disease-specific information, we then harmonized the disease terms present in the title and metadata of the datasets with the help of the Human Disease Ontology (DOID) (Schriml *et al*., 2018). Next, the disease terms from patient samples were mapped to DOID entities using ZOOMA (https://www.ebi.ac.uk/spot/zooma), enabling us, in some cases, to automatically find DOID matches. However, the majority of the terms did not contain a perfect match to a DOID entity so ZOOMA proposed the closest match. Based on this set of proposed DOID entities, we manually evaluated whether the term had been correctly assigned or a DOID entity that could more accurately represent the disease was available. Through this process, we also identified false positives terms which had not been successfully filtered in the previous steps. In these cases, the metadata did not contain sufficient information, though this information was present in the dataset title. Thus, using the title information, we removed such false positives terms following manual inspection.

To maximize the coverage, we conducted a final processing step where we intended to group similar diseases together under a common label. For that, we leverage the ontology network structure and visualize it as a hierarchical tree with a focus on selected branches (i.e., “immune system disease”, “nervous system disease”, and “cancer”). Next, we manually identify close neighbors for terms that have few samples in order to merge into a more general term that still accurately describes the original term. The veracity of the likeliness of the disease terms in the selected clusters to be used as a single gene expression set were verified by a clinician before re-mapping. **(Supplementary Text).**

After this final grouping step, we also filtered datasets to fulfill the following criteria: i) ensure every disease has a minimum of 50 samples to increase the stability of the co-expression network, ii) ensure a minimum of 2 datasets per disease, and iii) exclude samples with the “cancer” label as this term was too broad **(Figure 2, filter F).** Thus, we have 38,621 samples from 469 datasets as 63 distinct diseases and one control group **(Supplementary Table 1).** To facilitate the grouping of control samples, we first harmonized all samples coming from datasets used to generate the disease networks that correspond to controls by giving them a common label (i.e., “normal”) **(Figure 2, filter G)**. Applying the previously described filtering steps resulted in 35,025 samples from 323 datasets that were selected. Finally, not all datasets comprised the raw data required to generate the co-expression networks which are solely based on 279 datasets (20,748 samples) **(Figure 2, filter H).** The final list of datasets with their respective disease labels can be seen in **Supplementary Table 1** and can be visualized according to their DOID hierarchy in **Supplementary Figure 1**.

Scripts to retrieve and process the datasets from ArrayExpress are available at https://github.com/CoXPath/CoXPath. We have also provided comprehensive documentation to modify the filtering steps and add extensions to the scripts.

#### 2.1.2. Generating co-expression networks

For each disease, expression data could then be used to construct co-expression networks to represent relationships between genes in different diseases. Therefore, the raw .CEL-files of the expression datasets were downloaded, pre-processed, and merged. After merging the samples from different datasets, a batch correction via ComBat (Johnson *et al*., 2007) was applied to the data to remove the effect corresponding to individual datasets. Finally, the probes were mapped to genes. If multiple probes mapped to the same gene, the most variable probe was kept. In the special case of the normal network, we would like to note that only control samples that were present in the disease datasets were used **(Figure 2, filter G)**.

The actual co-expression datasets were then constructed with the WGCNA package in R (Langfelder *et al*., 2008). In contrast to most common approaches that construct and analyze modules of the network based on hierarchical clustering, here we relied only on the topological overlap matrix (TOM). For each disease, we defined its co-expression network as the top 1% highest similarity in the TOM, corresponding to 2,036,667 edges for each co-expression network.

### 2.2. Building a human protein-protein interactome network

To systematically compare disease-specific co-expression networks against pathway knowledge, we built an integrative network comprising information from a compendium of well-established databases. This interactome was comprised of tens of thousands of human protein-protein interactions from six databases including KEGG (Kanehisa *et al*., 2021), Reactome (Jassal *et al*., 2020), WikiPathways (Martens *et al*., 2020), BioGrid (Oughtred *et al*., 2019), IntAct (Orchard *et al*., 2014), and PathwayCommons (Rodchenkov *et al*., 2020). We would like to note that the first three of the six databases were harmonized through PathMe (Domingo-Fernández *et al*., 2019). Additionally, for each of the six databases, only proteins that belonged to pathways from MPath (Mubeen *et al*., 2019), an integrative resource that combines multiple databases and merges gene sets of equivalent pathways, were included in the interactome, thus ensuring that each protein in the network was minimally assigned to a single pathway. The use of MPath to annotate proteins to pathways facilitated both the generation of a larger network and the avoidance of redundant pathways.

The resulting human interactome network has a total of 8,601 nodes and 199,535 edges. Not surprisingly, the vast majority of the nodes in the interactome are protein-coding genes, as these genes are transcribed into functional proteins with essential roles in the biological processes represented in pathway databases **(Figure 3a).** Among the edges of the interactome, association relations are the most prevalent (~73%), while causal relations including, increase, decrease, regulate, and has_component relations constitute the remaining relation types **(Figure 3b)**.

**Figure 3.**
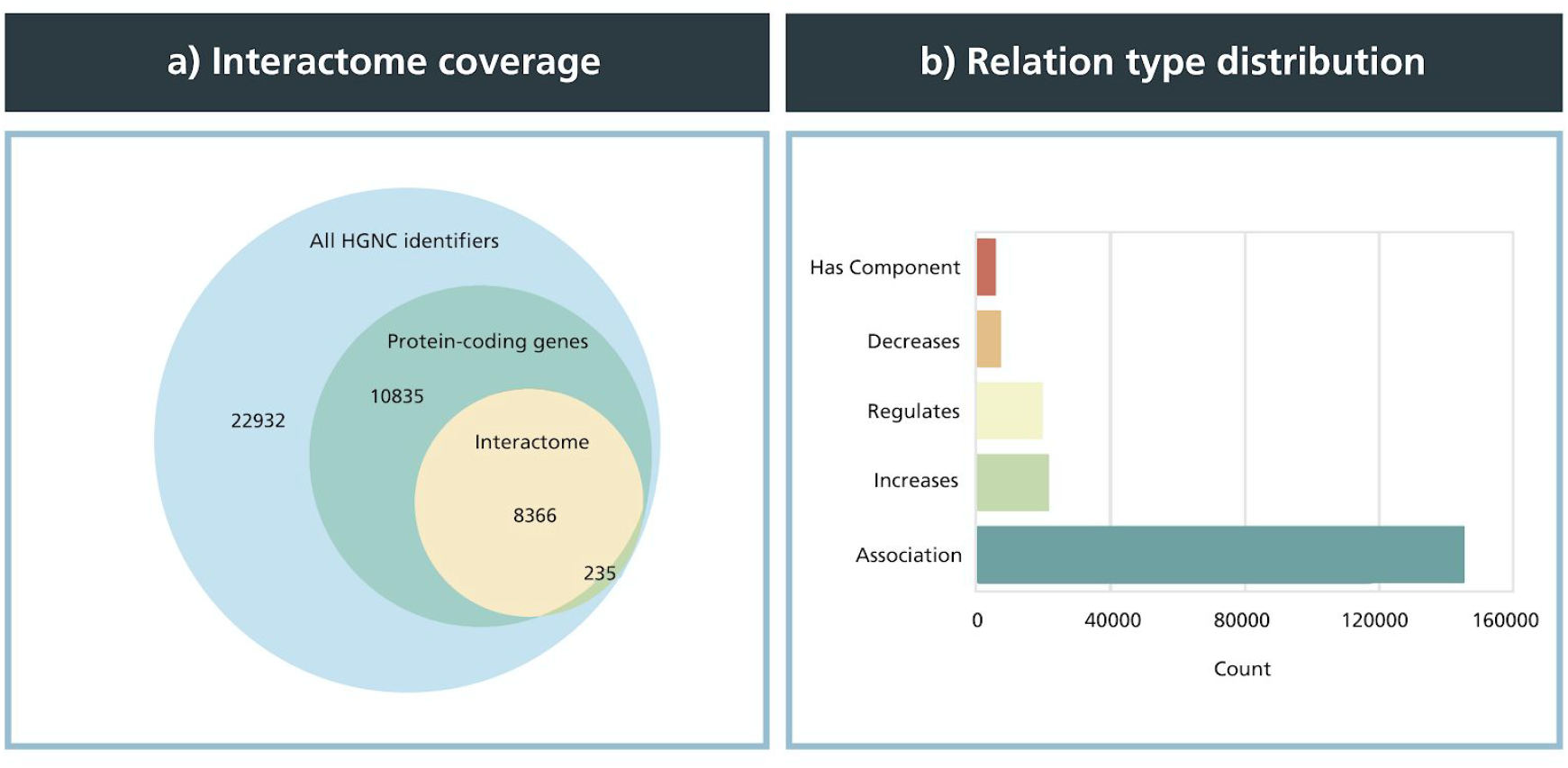
Node and edge type statistics of the human protein-protein interactome network. **a)** Venn diagram indicating the coverage of proteins in the interactome network with respect to all existing HGNC identifiers as well as protein-coding genes. The interactome contains ~8,600 unique HGNC identifiers, or 20% of the roughly 42,300 approved HGNC identifiers. In total, 97% of the HGNC identifiers of the interactome are protein-coding genes. **b)** Distribution of relation types in the interactome network. The largest proportion of relation types were associations, comprising nearly 73% of all ~200,000 edges, while causal relations, specifically decrease, regulate, and increase, made up ~25% of all relation types with roughly 50,000 edges.

### 2.3. Analyses

#### 2.3.1. Software and data used in network analysis and visualization

Network analyses were conducted using the methods and algorithms implemented in NetworkX (v2.5). KEGG pathways (Kanehisa *et al*., 2021) were downloaded on 03/08/2020 using ComPath (Domingo-Fernández *et al*., 2019). Network visualizations were done using WebGL, D3.js, and Threejs and the python-igraph package. The processed data and analyses are available at https://github.com/CoXPath/CoXPath.

#### 2.3.2. Meta-analysis of gene expression data

Differential expression analysis was performed using the Limma R package (Smyth, 2005) on the merged disease datasets described in subsection 2.1.1, which contained information on both patient samples and controls. This step yielded differentially expressed genes (DEGs) for 46 diseases in total from the original 63. DEGs for each disease were then filtered to include only those with an adjusted *p*-value < 0.05. DEGs across the 46 diseases were combined into a consensus by splitting the up- and down-regulated genes for each disease and taking the average adjusted *p*-values and log_2_ fold changes for all up- and down-regulated subsets separately.

#### 2.3.3. Quantifying the similarity between disease-specific co-expression networks and biological pathways

To investigate the consensus between the patterns present in each co-expression network and pathway knowledge, we superimposed each disease-specific co-expression network against pathways from KEGG and the interactome network described in subsection 2.2 using two different methods. Method 1 investigates every pairwise combination of nodes from the set of proteins *P* for a given pathway from KEGG (*C_P_*) to find the proportion of edges that exist in the disease co-expression network *D* = (*P*’, *E_D_*) between those node pairs, namely *edge overlap* (*edge overlap* = |{ ∀ *e_u,v_ s.t. u, v* ∈ *C_P_*; *u, v* ∈ *P’ and e_u,v_* ∈ *E_D_*}|) **(Equation 1)**. *P’* is the set of proteins in the co-expression network and *E* is the set of edges connecting the proteins.

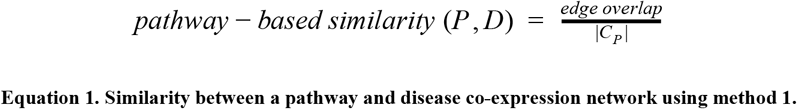

Similarly, applying a more stringent criterion to take into account the protein-protein interactome network, using a set of proteins *P* for a given pathway from KEGG, method 2 takes the interactome network *I* = (*U, E_i_*) and generates a subgraph *S* = (*V, E_s_*) containing only those nodes in *P* with edges in *E_i_* (with *V* = {*u*: *e_u,v_* ∈ *E_s_ and u,v* ∈ *P*∩*U*} *and E_s_* = {*e_u,v_*: *u, v* ∈ *P*; *u, v* ∈ *U*; *e_u, v_* ∈ *E_i_*}). Next, the proportion of edges on the interactome subgraph *S* that are also found in each disease co-expression network *D* = (*P*’, *E_D_*) are calculated **(Equation 2)**.

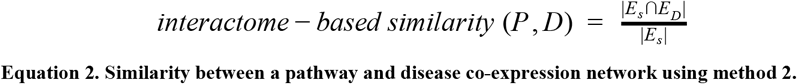

We would like to mention that we exclusively used pathway definitions (i.e., gene sets) from KEGG which contain a relatively fewer number of pathways in order to facilitate the interpretation of the analysis (e.g., Reactome contains over 2,000 pathways while KEGG has over 300). Nonetheless, in method 2, we overlay the KEGG gene sets onto the interactome network, ensuring that the analysis is not only restricted to biological interactions in KEGG.

#### 2.3.4. Pathway enrichment analysis

Overrepresentation analysis (ORA) was conducted employing a one-sided Fisher’s exact test (Fisher, 1992) for each of the pathways in KEGG (downloaded on 12-12-2020). A pathway is considered to be significantly enriched if its adjusted *p*-value is smaller than 0.05 after applying multiple hypothesis testing correction using the Benjamini–Yekutieli method (Benjamini and Yekutieli, 2001).

## 3. Results

In subsection 3.1, we first outline the diseases that fulfilled the criteria to generate the corresponding co-expression networks and investigate the characteristics of these networks. Then, in subsections 3.2 and 3.3, we analyze the disease-specific co-expression networks at the node- and edge-levels, respectively, while in 3.4, we compare the co-expression networks against pathway knowledge. Finally, in a case scenario (3.5), we demonstrate how a pathway-level analysis in a disease context can be leveraged to better understand the role of a specific pathway in a disease context.

### 3.1. Overview of disease-specific co-expression networks

From over 330 datasets that were categorized into distinct diseases, we systematically constructed 64 co-expression networks, 63 of which correspond to disease-specific co-expression networks, and the remaining corresponding to a control group co-expression network. **Figure 4A** summarizes the network size of each disease-specific co-expression network clustered by major disease indication for a total of ten disease categories and one unspecific group. Body system clusters (e.g., gastrointestinal system disease, immune system disease) were given priority for the classification of all cancers before considering the “other cancer” group. How each disease relates to its disease category cluster can be visualized on the DOID hierarchy in **Supplementary Figure 1**. The sarcoma co-expression network had the least number of nodes of all the networks (i.e., 5,450), while the ductal carcinoma in situ co-expression network had the highest number of nodes (i.e., 20,163). Generally, the networks within each disease category cluster tended to vary greatly in size. For example, the “immune system disease” category includes networks ranging in size from 5,754 to 18,449 nodes. Additionally, the number of co-expression networks within a disease cluster varied, with nearly half the disease groups containing between six and fifteen networks (i.e., gastrointestinal system disease, immune system disease, nervous system disease, respiratory system disease, and other cancer), while all remaining clusters contained less than five.

**Figure 4.**
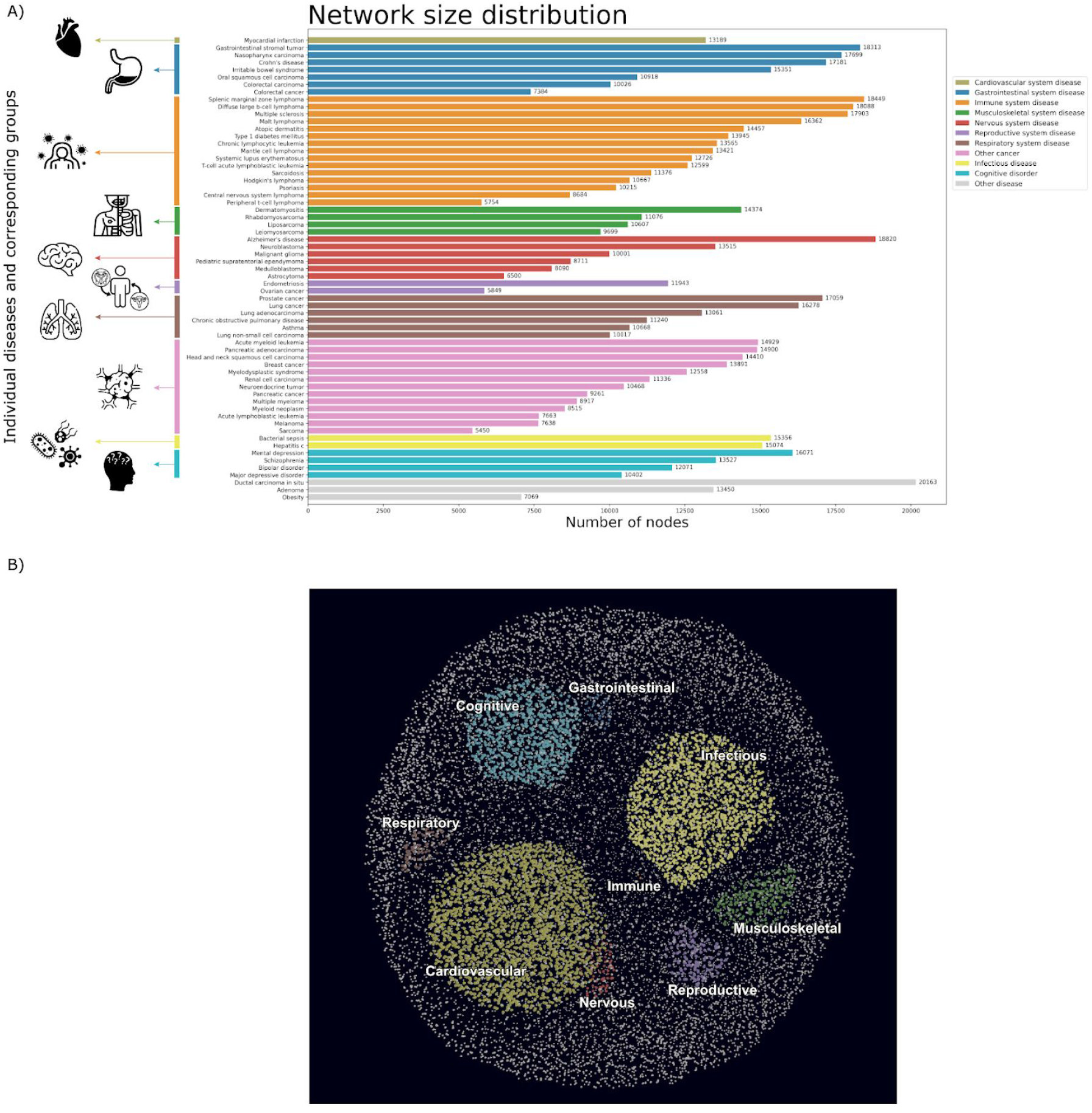
A) Overview of the size of each of the co-expression networks clustered by major disease groups. B) Merged co-expression network clustering proteins by their association to different disease groups. In A) each of the 63 diseases was grouped into one of ten categories (or a remaining leftover group). Here, we see the varied sizes of co-expression networks within their corresponding disease clusters. In B) association was determined by selecting the set of nodes which were present in all of the diseases of a given disease cluster (excluding “other” and “other cancer”), and eliminating those nodes which were also present in all diseases of other clusters. This resulted in unique sets of nodes which were guaranteed to be found in all diseases of the given cluster, but not in all of another cluster. As expected, we observed an inverse correlation between the number of diseases in a cluster and the size of the associated node subset. High quality versions are available at https://github.com/CoXPath/CoXPath/results/figures.

We also investigated whether a correlation exists between the number of samples or datasets used to create a co-expression network and the size of the network. No dependency of network size based on the amount of samples/datasets used was observed **(Supplementary Figure 2)**. The total number of datasets ranged from 1 to 27, while the total number of samples was between 9 and 2,515. The vast majority of disease co-expression networks were generated from 1 to 10 datasets and contained between 9 to 461 samples. We found that the resulting network size for each disease varied within a wide range (i.e., between ~6,000 and 20,000) and no discernible pattern was observed.

### 3.2. Investigating global trends of disease-specific co-expression networks at the node level

#### 3.2.1. Exploring the most and least common proteins of the co-expression networks

Here, we explored the most and least common proteins across all 63 disease-specific co-expression networks generated. We first identified the most common proteins as those that occur in the highest number of disease co-expression networks **(Supplementary Figure 3).** We found that none of the proteins were present in all co-expression networks as we were only interested in considering the top 1% strongest correlations in each network (i.e., the selected cut-off; see Methods subsection 2.1.2). On the other hand, TXLNGY and NCR2 were the most common proteins, occuring in 60 out of the 63 disease co-expression networks. Nonetheless, we were able to identify 48 proteins present in at least 57 of the 63 diseases.

We next investigated whether proteins in the disease co-expression networks could consistently be identified in our interactome network to infer how well these proteins have been studied and reported in the literature. We refer to proteins that could consistently be found across the majority of disease co-expression networks and were also present in the interactome as the most common proteins of the disease networks and the most highly connected proteins of the interactome. Surprisingly, we found that only 30-33.4% of the most common proteins (with cut-offs between 50 and 54 out of 63) of the disease co-expression networks were present in the interactome. Similarly, for an approximately proportional range of these most common disease proteins against the most connected proteins of the interactome (i.e., top 100-400 proteins), little to no overlap was observed **(Supplementary Figure 5)**. We also found that the average number of relations for the proteins in the interactome that overlapped with the approximately top 400 most common proteins in the disease networks (~33 relations) was lower than the average number of relations overall in the interactome (~46 relations). We then sought to verify whether the most common proteins in the disease co-expression networks could also be found in pathway databases, identifying only a small proportion (i.e., 29-31%) of these proteins in KEGG **(Supplementary Table 2)**. When comparisons were made against KEGG pathway annotations, we observed that these few most common proteins had, on average, a slightly lower number of pathway annotations (~14.8) than the average number of annotations for all proteins in the pathway database (~16). Taken together, these findings indicate that though these proteins are the most common across all disease co-expression networks, they tend to be underrepresented in the scientific literature.

Among the proteins in common between the top 400 most highly connected proteins of the interactome and the most common proteins in the disease co-expression networks, 13 proteins, including 3 members of the cytochrome P450 family of enzymes (i.e., CYP1A2, CYP2C9, and CYP3A4), a major ribosomal protein (i.e., RPL18), as well as key regulatory proteins such as CDK1, PRKCG, and PLCB2 were present in the overlap **(Supplementary Table 2 and Supplementary Figure 5)**. Similarly, we define proteins that could be consistently found across the majority of disease co-expression networks and were also present in KEGG pathway annotations as the most common proteins of the disease networks and the most common KEGG proteins. We examined the overlap between the top 400 most common disease proteins with the highest number of KEGG pathway annotations, and the most common proteins of the disease co-expression networks **(Supplementary Figure 8)**. Among the 22 proteins in common, we found 7 members of the human leukocyte antigens (HLA) system of proteins (HLA-B, HLA-C, HLA-DMA, HLA-DMB, HLA-DQB1, HLA-DRA, and HLA-G), as well as several proteins which were also in the overlap between the aforementioned most highly connected proteins of the interactome and most common proteins in the disease co-expression networks (i.e., CAMK2A, ELK1, GNAO1, and PLCB2) **(Supplementary Table 2 and Supplementary Figure 8)**.

Finally, we investigated the least common proteins in the disease co-expression networks and their overlap with those in both the interactome and pathway knowledge (**Supplementary Figure 6 and Supplementary Figure 9)**. Similar to the most common ones, we found that the majority of the least commonly occurring proteins in the co-expression networks were not present in the interactome nor in KEGG, suggesting that little is currently known of these proteins. Among the least commonly occurring proteins that overlapped with proteins from both KEGG and the interactome, we observed a significant number of proteins from the ZNF family (i.e., 42/54 (78%) from KEGG and 12/43 (28%) from the interactome overlap) **(Supplementary Table 2)**. This family is one of the largest protein families and is known to regulate a wide range of biological processes, while some of its members have already been associated with several disorders (Cassandri *et al*., 2017). Thus, it may be interesting to investigate proteins that are specific to a particular disease, or a few distinct diseases, in detail. As an example, we observed that TWIST1, one of the least commonly occurring proteins and a well-known oncogene (Gort *et al*., 2008), was exclusively present in only 25 diseases and over 50% of them were cancers **(Supplementary Table 2 and Supplementary Figure 10).**

#### 3.2.2. Meta-analysis on consistently differentially expressed genes across diseases

Differential gene expression analysis was performed in order to pinpoint genes which were consistently significantly differentially expressed between patient and control samples across 46 diseases. The average of all genes in these diseases that were up-regulated as well as the average of all genes that were down-regulated were independently calculated. **Figure 5** jointly reports the comparison of the negative log_10_ adjusted *p*-values versus log_2_ fold changes of all independently averaged up- and down-regulated DEGs in the 46 diseases. We found that nearly all genes were, to some degree, up-regulated in one or more diseases and down-regulated in at least one other, while only CCDC43, JADE3, RPL22L1, SOCS1, and TOR3A were exclusively up- and CAVIN2 and ZSCAN18 down-regulated across all diseases they were present in. In all, nearly 20,000 unique genes were significantly differentially expressed (adjusted *p*-value < 0.01), with ~17,600 up-regulated DEGs and ~15,600 down-regulated ones.

**Figure 5.**
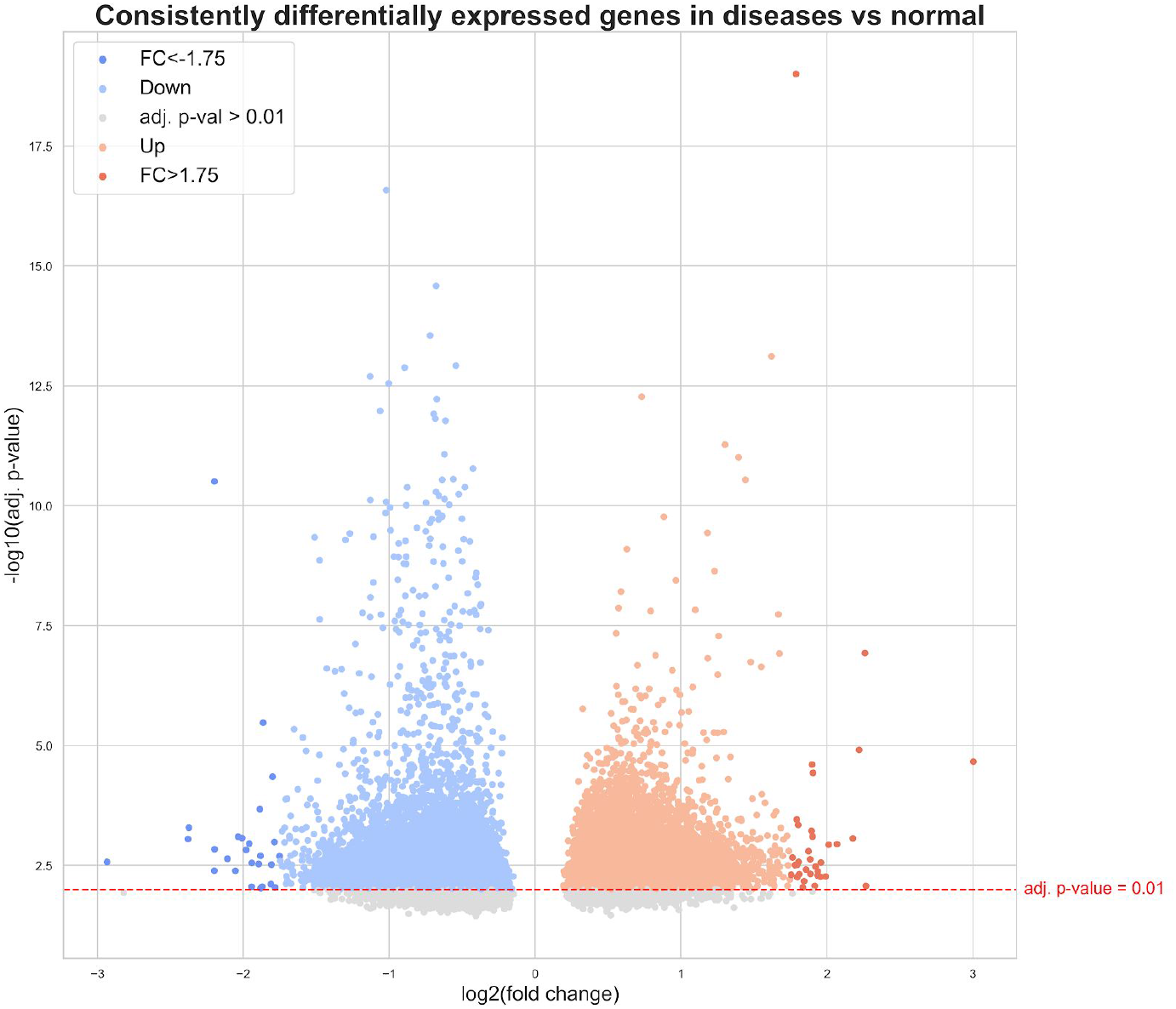
Consensus for consistently differentially expressed genes. Genes for 46 diseases were split into two subsets: those that were up-regulated and those that were down-regulated in that disease. The average consensus was taken for all up- and down-regulated subsets separately and is shown here. Nearly all genes could be found in both the up-regulated and down-regulated consensus as most genes are up-regulated in at least one disease as well as down-regulated in at least one other disease. In total, 19,666 unique genes were significantly up- or down-regulated (i.e., adjusted *p*-value < 0.01; above red line). Of the significantly differentially expressed genes, 17,643 genes were upregulated and 15,634 genes were downregulated. The most significantly differentially expressed genes were defined as additionally having a |*log_2_ fold change*| > 1.75, resulting in 26 most downregulated genes (dark blue) and 34 most upregulated genes (dark orange).

We then applied a |*log_2_ fold change* | threshold of 1.75 to identify significantly (adjusted *p*-value < 0.01) differentially expressed genes with the most extreme average log_2_ fold change values. This threshold was selected as it yielded a reasonable number of DEGs to investigate (i.e., 60), whereas more commonly used thresholds, such as |*log_2_ fold change*| > 1.5, yielded over 200. Among the genes that were found to be significantly differentially expressed at the extremes, 34 were the most up-regulated and 26 were the most down-regulated **(Supplementary Table 8)**.

These genes were then compared to the top 500 most and least common disease proteins. Of the genes that were the most up-regulated, CDK1 was also among the top 500 most common disease proteins, while CRNDE, DEPTOR, and RASD1 were among the 500 least common. Similarly, for genes that were the most down-regulated, only S100A8 was among the top 500 most common disease proteins while no genes overlapped with the 500 least common disease proteins. Additionally, we found that four of the most up-regulated genes belonged to the collagen group of protein (i.e., COL11A1, COL1A1, COL1A2, and COL3A1), while some protein families (i.e., S100 protein family, SLC, and SYNP) could be found both in the most up- and down-regulated genes.

Of the most significantly highly up- and down-regulated genes (i.e., adjusted *p*-value < 0.01; |*log_2_ fold change*| > 1.75), we examined their expression changes in each of the individual diseases they were involved in. Interestingly, we found a group of genes (i.e., AMPD1, BEX5, DEPTOR, IGF1, JCHAIN, MARC2, MTUS1, and NDFIP2) that were highly up-regulated in only two of nearly 20 diseases they were in (i.e., myeloid neoplasm and multiple myeloma, grouped in the other cancers cluster), and down-regulated in nearly all of the remaining. Thus, although these genes were down-regulated in the vast majority of the diseases they were involved in, they still appeared among the most significantly highly up-regulated genes due to their significantly high up-regulation in the two aforementioned cancers. This trend has been documented for DEPTOR, with low expression of the gene observed in most cancers, yet high overexpression seen in a group of multiple myelomas (Peterson *et al*., 2009). Similarly, among the genes with significantly high down-regulation, we identified a subset of genes (i.e., ASH1L-AS1, CXCL8, DUSP4, EPC1, PANK2, PCIF1, PHLDA1, and PMAIP1) that were only highly down-regulated in peripheral T-cell lymphoma, whilst being up-regulated in nearly all of the remaining diseases they were in (i.e., 17 diseases on average). This pattern has been identified with the overexpression of DUSP4, a tumor suppressor, in certain cancer types (Ratsada *et al*., 2020), whereas the loss of its expression caused by epigenetic dysregulation has been observed in at least one type of lymphoma (Schmid *et al*., 2015) Finally, the meta-analysis revealed that one gene with significantly high down-regulation, SLC8A1, was only significantly down-regulated in a group of nervous system diseases (i.e., medulloblastoma, pediatric supratentorial ependymoma, malignant glioma, astrocytoma, and to a lesser degree, Alzheimer’s disease), not altogether surprising as the SLC8 gene family of sodium-calcium exchangers, which includes SLC8A1, have been shown to play important regulatory roles in the control of central nervous system functions (Spencer *et al*., 2020). In contrast, SLC8A1 was only identified as having significantly high up-regulation in multiple myeloma.

### 3.3. Investigating global trends of disease-specific co-expression networks at the edge level

In this subsection, we explored the most common edges of the disease co-expression networks and compared them against the normal co-expression network and the interactome. We first assessed whether there were any edges specific to particular disease networks, identifying 57,774,118 unique edges in total (i.e., 45% of all edges). This was to be expected, as we exclusively focused on the 1% strongest correlations from the initial hundreds of millions of possible edges, which led to most of the edges in our resulting co-expression networks to be specific to a single disease. Although this unique, disease-specific set of edges are worth exploring, due to the considerably large number of edges in the co-expression networks, we restricted our analysis to the most common edges in the co-expression networks. We found that 21 edges were in more than 70% of the diseases (44/63) and 202 in more than 50% of the diseases (32/63). Interestingly, of those 21 edges that were in 70% of the diseases, we observed that 6 of the 13 proteins which are encoded by genes in the Y chromosome appeared in 5 edges each (i.e., RPS4Y1, USP9Y, DDX3Y, KDM5D, EIF1AY, and TXLNGY). Additionally, we found that nearly half of these 21 edges involved a protein of the Metallothionein family (i.e., MT1H, MT2A, MT1HL1, MT1X, and MT1G), involved in the regulation of transcription factors and in cancers (Gumulec *et al*., 2014).

The most common edges in the disease co-expression networks were then compared to the normal co-expression network to identify correlations between the two, assuming that proteins involved in these edges would have basal levels of expression and that they may not be relevant to a disease-specific context. Specifically, when the most common edges in the disease co-expression networks were compared to a proportionate range of edges with the strongest correlations in the normal network (i.e., from 1,000 to 10,000 edges), we found that between 19% and 17% of the edges consistently overlapped, respectively. Focusing on these ~19% of edges that were shared between the normal and most common disease networks, we were then interested in investigating whether these edges could also be found in the interactome, finding an overlap of only 8%. This number decreased to 4% as the number of edges being compared in disease against normal co-expression networks increased (i.e., between the top 1,000 and 10,000 most common edges). Additionally, from these 8% to 4% of edges which overlapped with the interactome, we looked at the top 10 most connected proteins, consistently identifying the same proteins as the number of edges in the comparison increased. Furthermore, we found that the direct overlap between the top 1,000 most common edges of the disease networks with the interactome was only 4%, while the overlap between the interactome and the top 1,000 most common edges of the disease networks which were not among the top edges of the normal network was 2%. Because this latter group of edges represents the top edges of the disease co-expression networks (but not of the normal) which overlap with the interactome, they may also warrant further investigation as they are more likely to consistently appear across diseases than in normal networks.

### 3.4. Overlaying co-expression networks with pathway knowledge supports the identification of disease associated pathways

In this subsection, we systematically overlayed pathway knowledge with disease co-expression networks to reveal the consensus and/or differences between the latter group of networks and well-established protein-protein interactions in pathway databases. Given that strongly co-expressed genes can be used as a proxy for functional similarity (Paci *et al*., 2021), it can be inferred that genes that are co-expressed could also be involved in the same pathway. In other words, we assume that if a given pathway is relevant to a disease, the proteins in the pathway would be strongly correlated in the disease co-expression network. Thus, following this assumption, we were interested in identifying the pathways associated with each of the investigated diseases. Using pathways from KEGG, we applied two methods which, i) map pathway knowledge to disease co-expression networks and ii) map pathway knowledge to the interactome, and the mapped portion of the interactome to disease co-expression networks **(see Methods).**

As expected, we noted that the results of both methods were nearly identical, indicating that pathway proteins were readily mappable to the interactome. Nonetheless, we found that the second method resulted in generally higher similarity values as it only considered edges that were identifiable in the interactome, rather than edges resulting from all possible combinations of pathway proteins (**Supplementary Figure 10**). Overall, clearly noticeable patterns were discernible, with groups of pathways showing variable levels of similarity in specific diseases and disease clusters (**Figure 6**).

**Figure 6.**
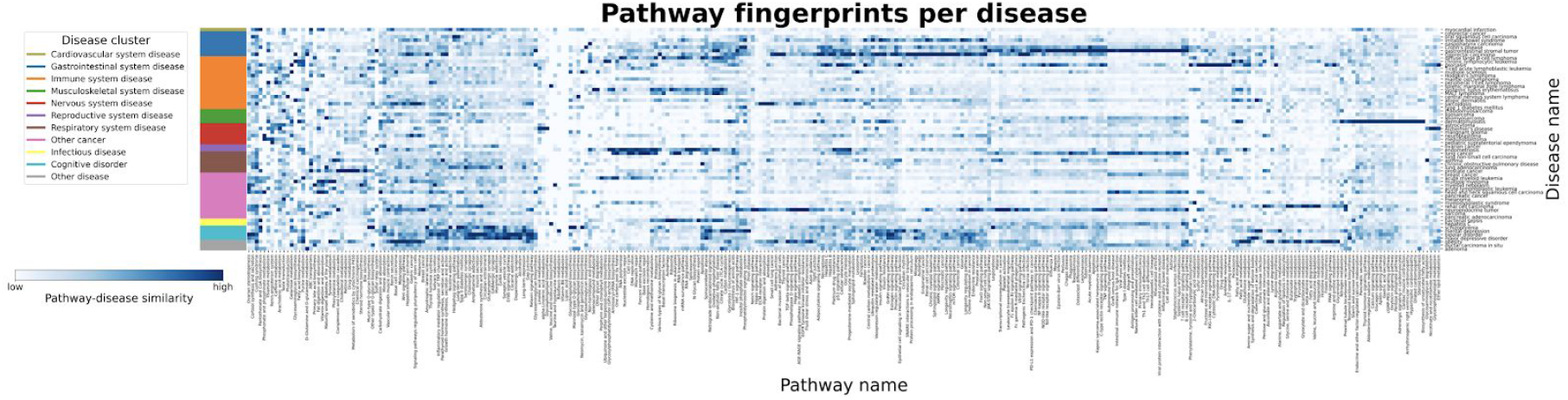
Mapping disease-specific expression patterns with pathway knowledge via network similarity. The heatmap illustrates the consensus similarity between KEGG pathways and disease co-expression networks. Similarity was defined as the percent of neighbors existing in a disease co-expression network out of all possible pairs of proteins from KEGG pathways (i.e., pathway-disease similarity), with lighter values corresponding to a lower similarity and darker values corresponding to higher similarity. The values (given as the percent of neighbors found) were standardized to a feature range from 0-1 for each pathway and pathways with similar values were grouped together. To ease the identification of patterns of pathway fingerprints across similar diseases, diseases were grouped by the previously defined clusters **(Figure 4)**. A high quality version of this figure is available at https://github.com/CoXPath/CoXPath/results/figures.

In particular, we observed multiple diseases/disease clusters with higher similarity values for pathways relevant to the given disease/cluster. Among these clusters, a large group of pathways showed a high degree of similarity to cognitive disorders **(Figure 6; teal)**, including pathways for long-term potentiation, multiple neurotransmitter systems (i.e., serotonergic synapse, glutamatergic synapse, and dopaminergic synapse), long-term depression, alcoholism, and pathways for addictions (i.e., nicotine addiction, amphetamine addiction, morphine addiction, and cocaine addiction) (**Supplementary Table 3**). Not surprisingly, the pathway for long-term depression showed the highest similarity with the co-expression network for mental depression. Furthermore, the gastrointestinal system disease cluster **(Figure 6; blue)** contained co-expression networks with the highest level of similarity with several pathways, e.g. the pathways responsible for renal cell carcinoma, colorectal cancer, pathogenic *Escherichia coli* infection, intestinal immune network for IgA production, and inflammatory bowel disease (**Supplementary Table 4**). Additionally, a broad group of pathways showed the highest similarity values for the two reproductive system diseases (i.e., endometriosis and ovarian cancer) **(Figure 6; purple)** over all other diseases and disease clusters **(Supplementary Table 5)**. Interestingly, we found that several cancers, including gastrointestinal stromal tumor, lung cancer, head and neck squamous cell carcinoma, neuroendocrine tumor, hepatitis C, breast cancer, and ductal carcinoma in situ shared a common pattern of similar pathways (**Supplementary Table 6**). Among the diseases, dermatomyositis was particularly distinguishable above all others, displaying notably higher similarity to several pathways **(Supplementary Table 7)**.

Altogether, we have demonstrated how by overlapping pathway knowledge to disease-specific co-expression networks, we can identify pathways associated with a particular disease. Additionally, we have also shown how this approach can be used to cluster diseases by the pathways they have in common, pointing to sets of potentially shared mechanisms across diseases.

### 3.5. Case scenario: in-depth investigation of the long term potentiation pathway in the context of schizophrenia

Previously, in subsection 3.4, we identified disease-associated pathways by calculating similarity between pathway knowledge and disease co-expression networks. To understand the mechanisms that underlie the similarity of a pathway to a given disease, in a case scenario, we next investigated the long term potentiation (LTP) pathway which had yielded high similarity to the schizophrenia co-expression network. An association between this pathway and schizophrenia has already been reported in the literature, with evidence indicating impairment of LTP in the disorder (Frantseva *et al*., 2008; Hasan *et al*., 2011).

The LTP pathway is categorized as a nervous system pathway in KEGG, with 35 edges between a set of 25 proteins/protein complexes **(Figure 7)**. As 19 of the nodes are protein complexes containing multiple proteins, the pathway covers a total of 67 unique proteins. By overlaying the co-expression network for schizophrenia with this pathway, we identified four major edges in common, all of which were well-established interactions within this particular pathway and formed a subgraph. These edges were among the most essential of the LTP pathway; interactions between protein kinase A and the NMDA receptor, Ca2+/calmodulin-dependent protein kinase II (CAMKII) and calmodulin, and the subsequent activation of AMPAR (Kristensen *et al*., 2011) and metabotropic glutamate receptors (Foster *et al*., 2018) by CAMKII play key roles in determining the strength of synaptic transmission and ultimately the expression of LTP (Herring and Nicoll, 2016).

**Figure 7.**
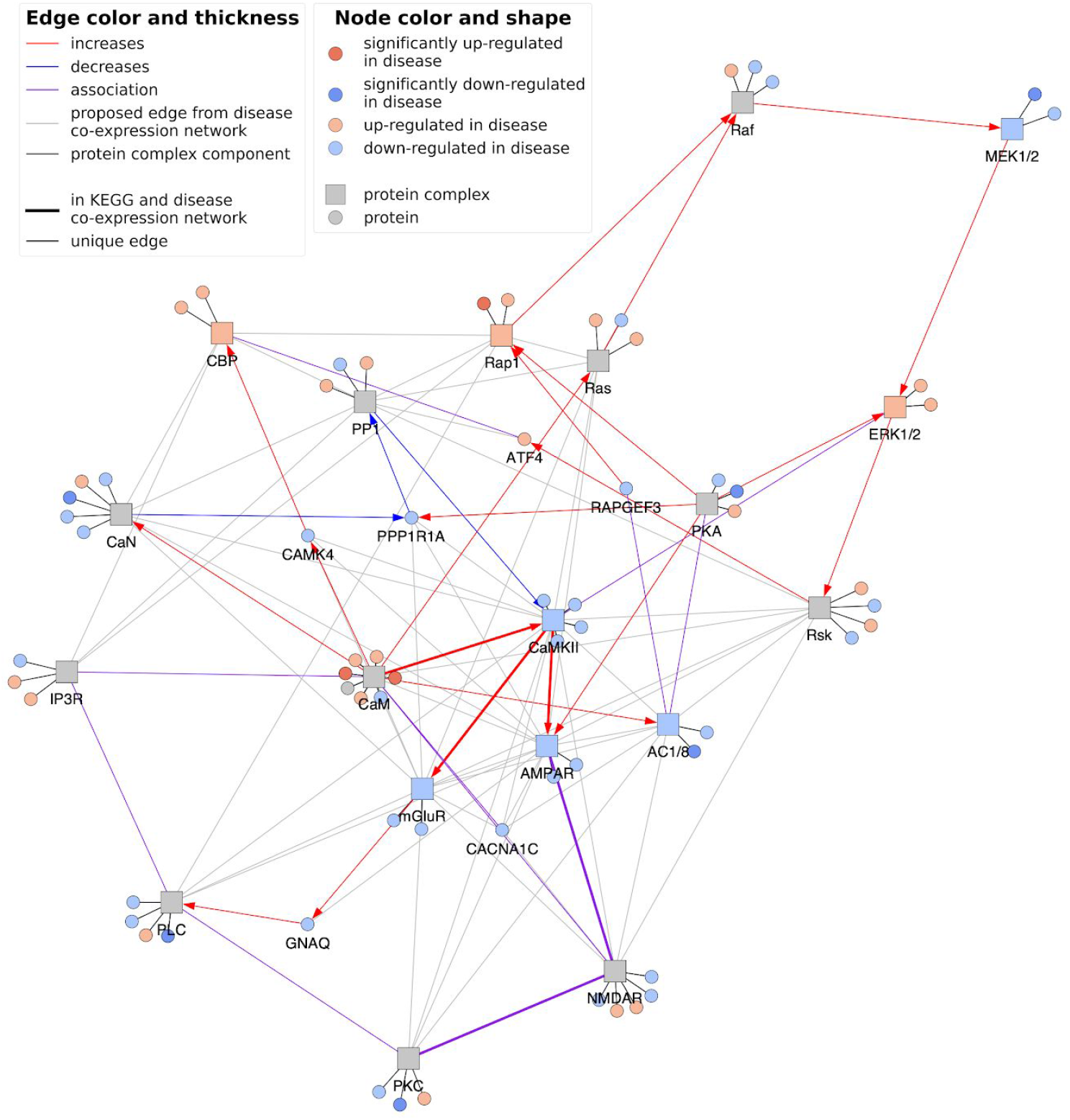
Long term potentiation (LTP) pathway in the context of schizophrenia. The figure depicts the overlap of the LTP pathway with the schizophrenia co-expression network. Protein-protein interactions and associations between proteins and/or protein complexes are displayed as colored edges, while black edges denote membership of proteins to protein complexes. Edges that were common to both the LTP pathway and the disease co-expression network are bolded, while grey edges denote correlations exclusively from the schizophrenia co-expression network. Differential gene expression analysis was performed and genes that were up- and down-regulated are colored orange and blue, respectively, with those that were significantly differentially expressed (i.e., adj. *p* value < 0.05) given less transparency. Protein complex nodes are then additionally colored if all members are in agreement with the direction of regulation. The code to generate this figure for any combination of disease co-expression network and pathway can be found at https://github.com/CoXPath/CoXPath/analysis/3.5_analysis.ipynb.

Interestingly, by overlaying the schizophrenia co-expression network with the LTP pathway, we found 53 unique correlations between proteins of the LTP pathway, indicating that the vast majority of proteins in this pathway were correlated in the co-expression network (**Figure 7**; grey edge), and demonstrating that indeed, proteins that are correlated in a given co-expression network can also be involved in the same biological process (Vella *et al*., 2017). 19 of these correlations were between calcium voltage channel complexes or calmodulin, which both have roles in the initial activation of the pathway, and other proteins (e.g., glutamate receptors). Similarly, there were approximately 20 correlations between all glutamate receptors present in the pathway and other proteins. The remaining correlations involved Erk/MAP kinase and cAMP, which ultimately regulate EP300 and CREBBP (which form the CREB binding protein complex) as well as ATF4. ATF4 is a transcription factor with multiple regulatory functions and whose polymorphisms have been associated with schizophrenia in male patients (Qu *et al*., 2008).

Lastly, we attempted to pinpoint candidate downstream pathways of LTP in the context of schizophrenia by investigating the edges of ATF4 given its role as a key regulator of the LTP pathway (Pasini *et al*., 2015). As ATF4 is strongly correlated with 70 other proteins in the co-expression networks, we conducted a pathway enrichment analysis as a proxy to reveal pathway crosstalks mediated by this protein **(see Methods)**. This analysis pinpointed four pathways from which three were involved in protein and RNA processing (i.e., ubiquitin mediated proteolysis, RNA transport, spliceosome), biological processes which have been linked with schizophrenia (McInnes and Lauriat, 2006; Glatt *et al*., 2011), while the fourth pathway, cell cycle, has also been associated with the disease (Fan *et al*., 2012; Katsel *et al*., 2008) (**Supplementary Table 9**). These findings indicate that there may be crosstalk between these pathways that could be explored in the future.

## 4. Discussion

Here, we have presented a systematic network-based approach that builds a bridge between disease signatures and pathway knowledge to better understand human pathophysiology. Our analysis has enabled us to globally evaluate the consensus between disease-specific transcriptomic data and an integrative human interactome network. Leveraging hundreds of transcriptomic datasets from over 60 major indications, we have explored the expression patterns observed in their corresponding co-expression networks at three different scales (i.e., at the node, edge and pathway levels). At each of these scales, we have investigated which proteins, subgraphs, and pathways could be associated with both disease-specific and shared mechanisms. Finally, we have presented a case scenario where we demonstrated how our approach can be used to investigate the role of a specific pathway in a disease-specific context.

There exist several limitations to this study. Firstly, we sought to improve the quality of the data by systematically integrating transcriptomic datasets from the same disease group, however, in doing so, we assumed that these datasets were equivalent. Although we attempted to address this assumption by enforcing a conservative inclusion and exclusion criteria as well as extensively curating the metadata associated with each dataset to group datasets into distinct diseases, disease heterogeneity for patients cannot be ignored. Secondly, we restricted this study to the most used platform in ArrayExpress in order to avoid possible effects caused by the array type, thus limiting the number of datasets that could potentially be used. Thirdly, since the cut-off chosen to generate the co-expression networks influences the resulting network (Yip and Horvath, 2007), we exclusively focused on the 1% strongest correlations. While this cut-off was well-suited for our large-scale approach, in the future, less restrictive cut-offs could be used to generate co-expression networks as well as other methods. For instance, Pardo-Diaz *et al*. (2021) recently presented a novel method that adds directionality into the co-expression network. Finally, while we constructed a human interactome network from multiple pathway and interaction databases, the majority of proteins from the co-expression networks could not be mapped to the network, highlighting the incompleteness of the current interactome.

Although we have demonstrated a proof-of-concept of our methodology across hundreds of datasets and in over sixty indications, we were only able to scratch the surface of the possible analyses that could be conducted with the resources generated within the context of this work. Thus, we have made the datasets and scripts generated in this study public to allow other researchers to conduct additional analyses on them. In the following, we outline several future applications and extensions of this work. Firstly, while we employed data from microarray technologies, the presented analysis could be expanded and/or validated by incorporating datasets generated from other platforms and technologies (e.g., RNASeq) or deposited in other databases such as GEO (Edgar *et al*., 2002) which, in turn, can facilitate the discovery of novel genes as well as allow us to add new indications and validate the current mechanisms identified in our analysis, respectively. However, conducting such an analysis would require extensive harmonization efforts at both the data and metadata level given the differences across chips and technologies, and the lack of structured metadata present in transcriptomic experiments. Secondly, the disease-specific co-expression networks generated in this work could be compared against well-established databases such as DisGeNet (Piñero *et al*., 2016) and OMIM (Hamosh *et al*., 2005) to propose novel gene-disease associations that can be integrated into these resources. Thirdly, other advanced network analysis methods could be conducted to analyze specific network motifs in the future. Fourthly, with prior enrichment of the presented networks with drug-target information, network-based drug discovery methods can be applied to identify candidate drugs and druggable pathways for the particular disease condition(s) (Peyvandipour *et al*., 2018; Rivas-Barragan *et al*., 2020). Finally, another potential line of research would be to apply our methodology on datasets generated from a variety of cell lines to identify cell-specific transcriptional patterns.

## Authors’ Contributions

MHA conceived the original idea. DDF designed and supervised the study with assistance from SM. TR implemented the pipeline to download, process and categorize the gene expression datasets. RQF and TR generated the co-expression networks for each group. SM and DDF generated the interactome network. RQF performed the analyses. RQF, TR, SM and DDF interpreted the results. ATK, MHA, and DDF acquired the funding. RQF, TR, SM and DDF wrote the manuscript.

All authors have read and approved the final manuscript.

## Acknowledgements

We would like to thank Sumit Madan for his help running the ZOOMA queries, Carlos Bobis-Álvarez for his assistance grouping similar indications, Chris W. Diana, Helena Hermanowski, and Carina Steinborn for their assistance generating the figures, and Daniel Rivas-Barragan for his valuable feedback.

## Funding

This work was developed in the Fraunhofer Cluster of Excellence “Cognitive Internet Technologies”. This work is supported by the German Federal Ministry of Education and Research (BMBF, grant 01ZX1904C).

## Conflict of Interest

DDF received salary from Enveda Biosciences.

## Notes

https://github.com/CoXPath/CoXPath

https://doi.org/10.5281/zenodo.4572853

