## Supplementary File for "Towards a global investigation of transcriptomic signatures through co-expression networks and pathway knowledge for the identification of disease mechanisms"

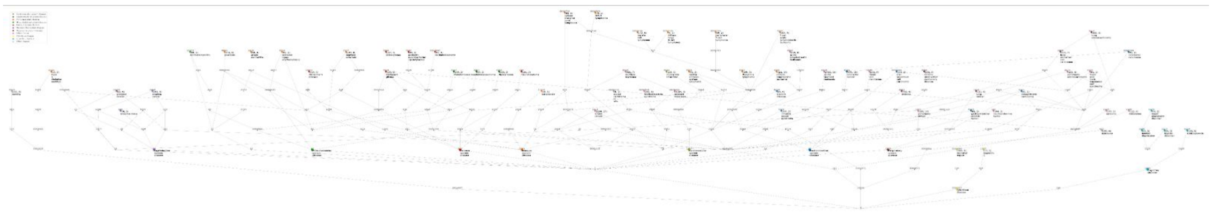

**Supplementary Figure 1.** Tree showing where in the hierarchy the groups lie in regards to each other according to the Human Disease Ontology (DOID). All diseases have DOID labels, those for which we have a disease co-expression network additionally have their disease name, tuple with (number of samples, number of datasets used), and are colored according to cluster assignments. Cluster assignment roots are also labelled and colored. A high quality version of this figure is available at https://github.com/CoXPath/CoXPath/results/figures.

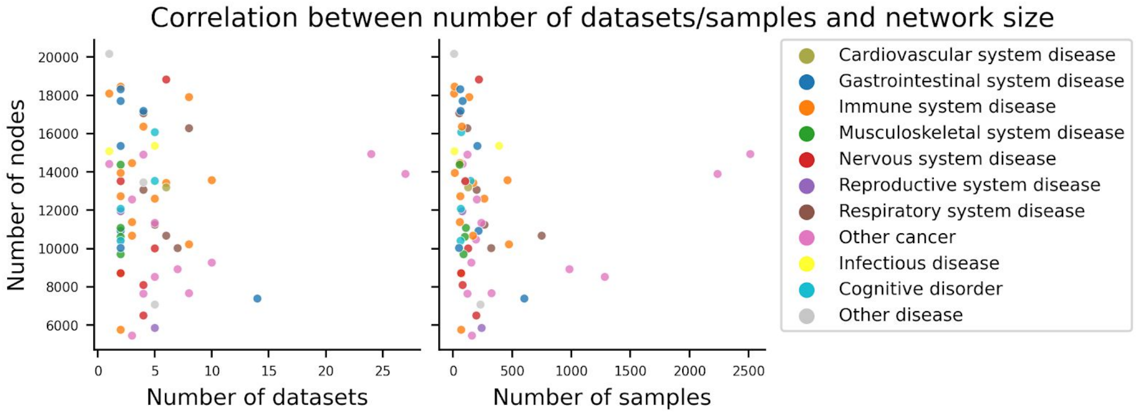


**Supplementary Figure 2. Scatterplot illustrating the correlation between number of nodes to number of datasets (left) and number of samples (right) for all diseases.** Each of the 63 disease-specific co-expression networks are shown as points on both plots and colored by their assigned clusters. Neither the number of datasets nor number of samples used to create the co-expression networks show any correlation to the resulting size of the network.

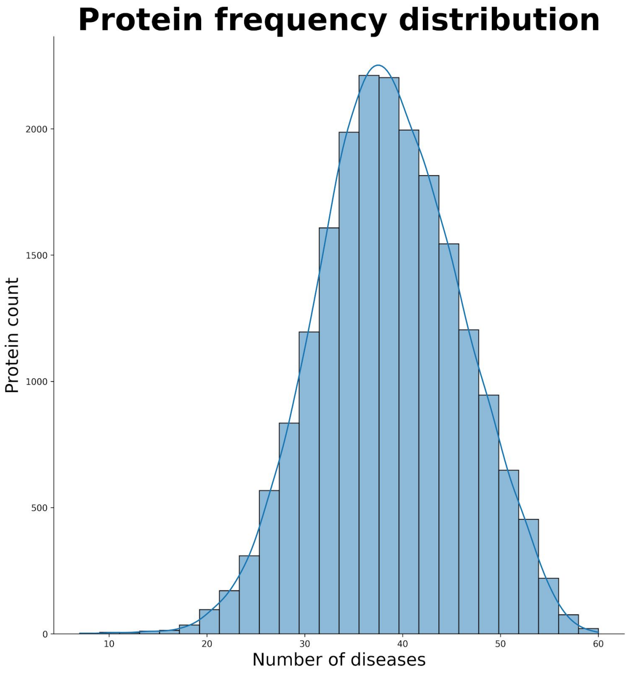


**Supplementary Figure 3. Distribution of the frequency of all proteins across the disease co-expression networks.** All proteins from all disease co-expression networks were collected to find how many diseases each protein occurs in. The resulting histogram shows how many proteins occur in any given number of the diseases. No protein was present in more than 60 diseases. The majority of the proteins are present in ~35-40 of the diseases.

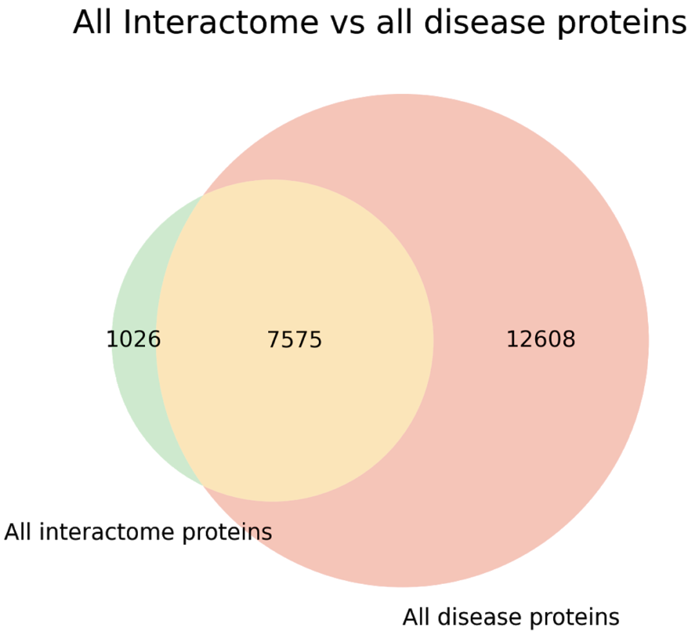


**Supplementary Figure 4. Venn diagram of the overlap between all proteins of the interactome and all proteins of the disease co-expression networks.**

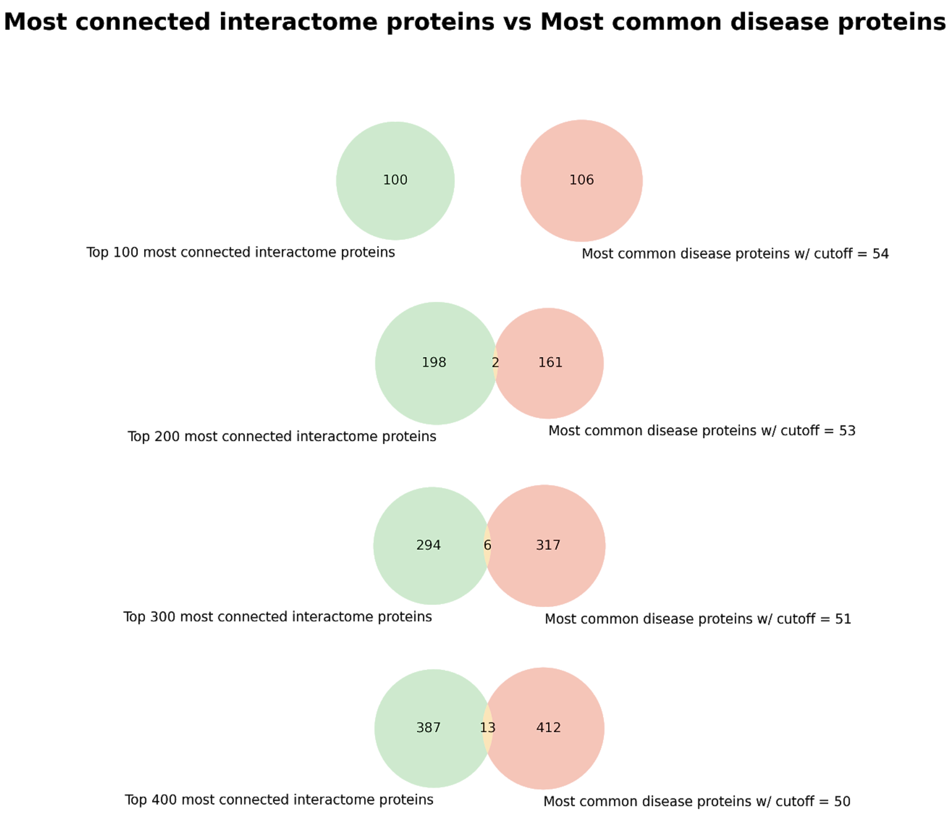


**Supplementary Figure 5. Venn diagram of the overlap between the most well-connected proteins of the two network types.** From the intersection of all interactome and disease proteins (i.e., **Supplementary Figure 4**), those of which are the most highly connected in the interactome (green) and most common in the majority of disease co-expression networks (red), are compared at varying levels (i.e., ~100-400 proteins). As the number of the most highly-connected proteins of the interactome increases relatively proportionately with the number of common proteins in the disease co-expression networks, little overlap is observed.

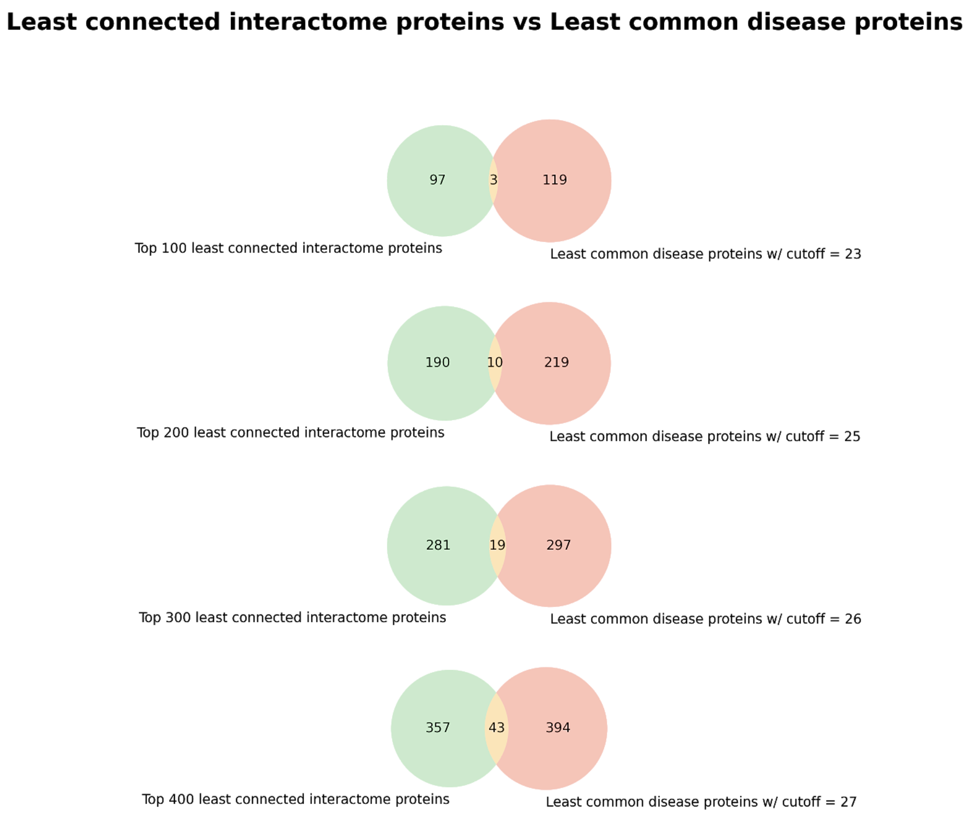


**Supplementary Figure 6. Venn diagram of the overlap between the least connected proteins of the two network types.** From the intersection of all interactome and disease proteins (i.e., **Supplementary Figure 4**), those of which are the least connected in the interactome (green) and least common in the majority of disease co-expression networks (red), are compared at varying levels (i.e., ~100-400 proteins). As the number of the least connected proteins of the interactome increases relatively proportionately with the number of least common proteins in the disease co-expression networks, some overlap can be observed.

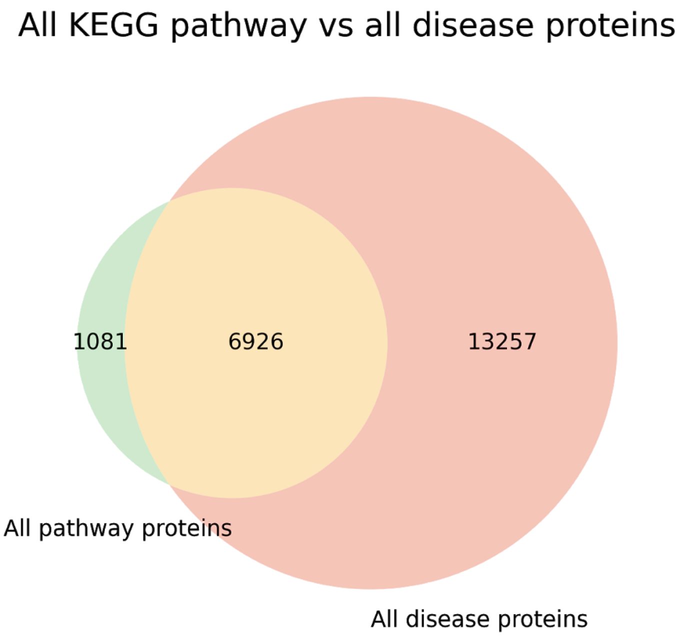


**Supplementary Figure 7. Venn diagram of the overlap between KEGG pathway proteins and all proteins of the disease co-expression networks.**

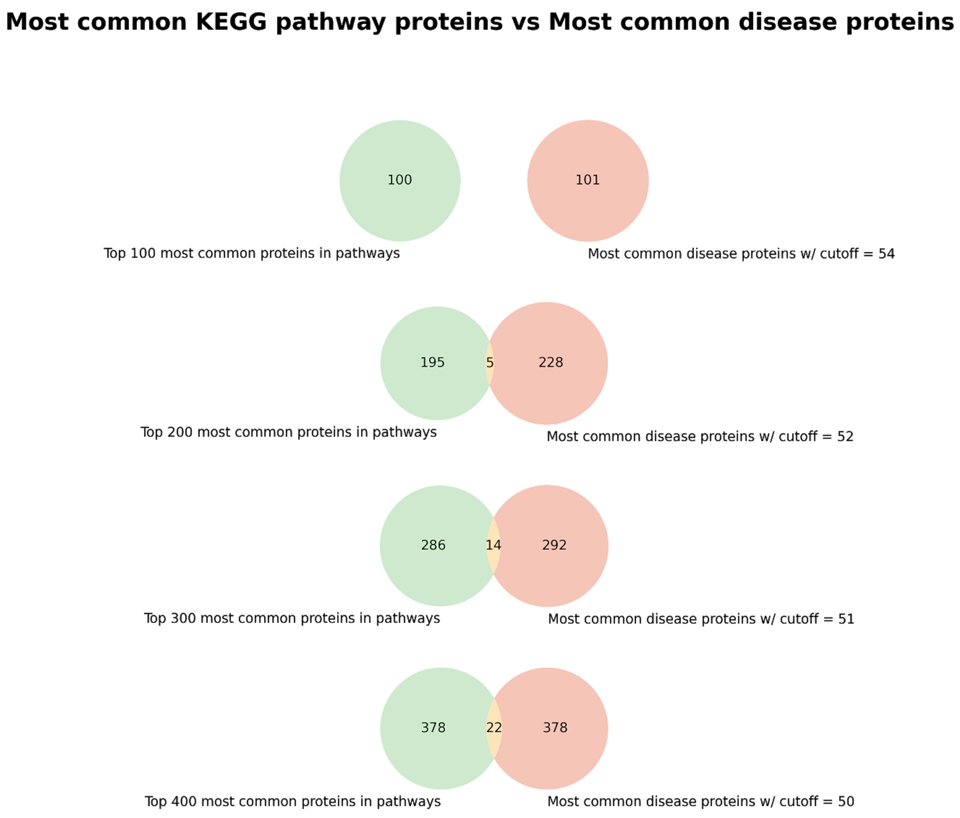


**Supplementary Figure 8. Venn diagram of the overlap between the most common proteins from KEGG and the disease co-expression networks.** From the intersection of all KEGG pathway and disease proteins (i.e., **Supplementary Figure 7**), those of which are the most common from KEGG (green) and most common in the majority of disease co-expression networks (red), are compared at varying levels (i.e., ~100-400 proteins). As the number of the most common KEGG pathway proteins increases relatively proportionately with the number of most common proteins in the disease co-expression networks, little overlap is observed.

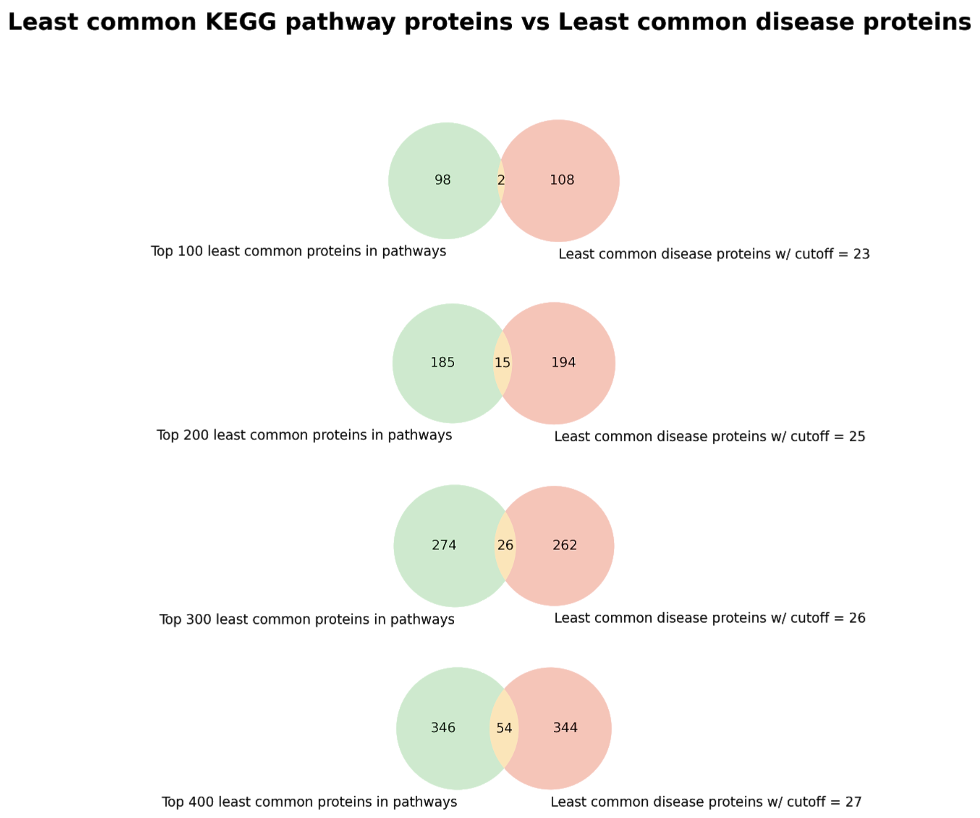


**Supplementary Figure 9. Venn diagram of the overlap between the least common proteins from KEGG and the disease co-expression networks.** From the intersection of all KEGG pathway and disease proteins (i.e., **Supplementary Figure 7**), those of which are the least common from KEGG (green) and least common in the majority of disease co-expression networks (red), are compared at varying levels (i.e., ~100-400 proteins). As the number of the least common KEGG pathway proteins increases relatively proportionately with the number of least common proteins in the disease co-expression networks, some overlap can be observed.

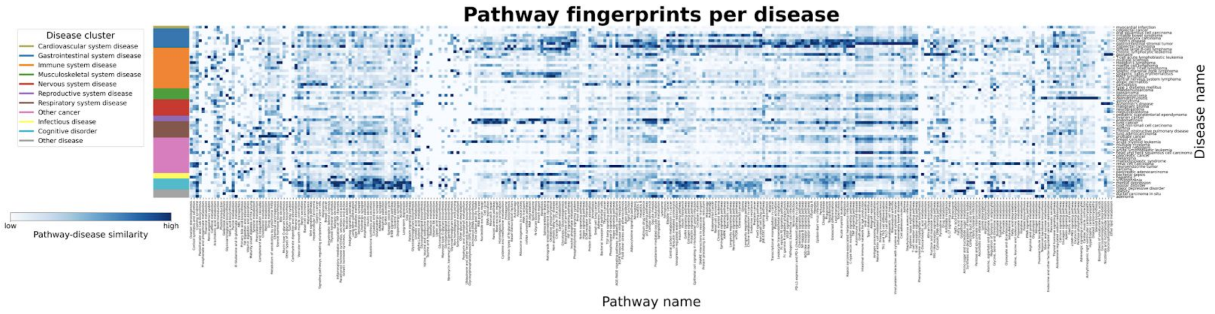


**Supplementary Figure 10. Mapping disease-specific expression patterns with pathway knowledge via network similarity.** The heatmap illustrates the consensus similarity between KEGG pathways and disease co-expression networks. Similarity was defined as the percent of existing edges in a disease co-expression network from the interactome subset with only proteins belonging to the given pathway, with lighter values corresponding to a lower similarity and darker values corresponding to higher similarity (i.e., pathway-disease similarity). The values (given as the percent of edges found) were standardized to a feature range from 0-1 for each pathway and pathways with similar values were grouped together. To ease the identification of patterns of pathway fingerprints across similar diseases, diseases were grouped by the previously defined clusters **(Figure 4)**. A high quality version of this figure is available at https://github.com/CoXPath/CoXPath/results/figures.

**Supplementary Text 1. Manual curation of disease group clusters**

Some of the groups from the mapped terms have too few samples to build co-expression networks from, however many terms appear to be similar enough that they can be considered one. E.g., 'chromophobe renal cell carcinoma', 'papillary renal cell carcinoma', and 'clear cell renal cell carcinoma', which individually wouldn't have enough data to create a co-expression network, could be re-mapped under 'renal cell carcinoma'. Changes like these are explored in this additional manual curation step.

The hierarchy of the disease ontology was visualized as a tree, with nodes from which we have samples of, highlighted and annotated. Branches that include no highlighted nodes are left out of the visualization, and every terminal node is a highlighted one (no need to show all child terms) to simplify the tree. Three important branches were selected for manual inspection and curation, ones known to be included in many datasets. These groups are "immune system disease", "nervous system disease", and "cancer".

Highlighted nodes were manually inspected in these three groups to identify close neighbors and determine if biologically similar enough to merge. Clusters were determined and marked for merging, with a maximum of a two-node jump with the exception of three-node jumps for breast carcinoma. These potential clusters were discussed in detail with a medical doctor to confirm that the diseases in them were biomedically similar enough to combine gene expression data as one. Those verified clusters were re-mapped to the selected broader term.

With the newly re-mapped data, a few more filters were used before selecting the final groups to be used. These were: the group should contain at least 50 samples (to ensure enough for a better co-expression network), the group should come from at least two datasets (to thin out the result), and excluding those mapped simply as 'cancer' (this term is too general to provide significant results despite making it past the first filters).
